# *Hypsugo stubbei* sp. nov., a novel cryptic bat species of the genus *Hypsugo* (Vespertilionidae, Chiroptera, Mammalia) from Mongolia. Results of the Mongolian-German Biological Expeditions Since 1962, No. 347.

**DOI:** 10.1101/2021.03.30.437544

**Authors:** D. Dolch, M. Stubbe, N. Batsaikhan, A. Stubbe, D. Steinhauser

## Abstract

The occurrence of two members of the genus *Hypsugo*, namely *H. alaschanicus* and *H. savii caucasicus*, have been reported for Mongolia in the literature. Due to various taxonomic reassignments within and between genera, the number of records for the genus *Hypsugo* in Mongolia is quite scarce and sometimes not resolved at species or subspecies level. Despite recognition of the two above-mentioned species, recent reports based on genetic analyses describe only new and further records of *H. alaschanicus*. Thus, it exists a large uncertainty regarding the occurrence and distribution of *H. savii caucasicus* in Mongolia. Here, our efforts in gaining a deeper understanding towards the occurrence and distribution of *Hypsugo* species in Mongolia are described.

A combination of genetic and morphological analyses of collected material from *Hypsugo* specimens revealed the existence of a genetically largely distant *Hypsugo* clade. Therefore, a new and cryptic *Hypsugo* species is proposed which is named after Prof. Dr. Michael Stubbe for his continuous, long-standing and significant contributions into the biological exploration of Mongolia. *Hypsugo stubbei* sp. nov. differs by at least 8.4 % and 9 % to the closest Western Palearctic distributed *H*. cf. *darwinii* and *H. savii* as well as at least 11.3 % to the Easter Palearctic (including Mongolia) distributed *H. alaschanicus* based on the first 798 nucleotides of the gene encoding the mitochondrial ND1 (subunit one of NADH dehydrogenase). Neither a close proximity species based on the gene encoding the mitochondrial COI (cytochrome oxidase subunit one) could be found in publicly accessible nucleotide databases. While the cryptic *H. stubbei* sp. nov. reveals no obvious cranial and morphological differences, few external characteristics are dissimilar to both *H. alaschanicus* and *H. savii* (*caucasicus*). Currently, *Hypsugo stubbei* sp. nov. was found at four different locations in Mongolia. Among the 11 specimens captured, six facilitated a genetic assignment. Based on the current scarce data records, the species seems to occur mainly in the far west of Mongolia inhabiting semi-deserts and steppes up to high mountain areas. An overlapping distribution with *H. alaschanicus* cannot be excluded based on the limited data currently available.

## Introduction

For a long time the genus *Hypsugo* was grouped together with the actual genera *Vespertilio, Eptesicus*, and *Pipistrellus* into the *Vespertilioninae* prior separating this taxon group and assigning *Hypsugo* into the genus *Pipistrellus* (HILL & HARISSON 1987; PAVLINOV et al. 1995; KOOPMAN 1994). After further separation and delimitation of the “true” *Pipistrellus* (HORÁČEK & HANÁK 1986), karyology studies and genetic characteristics (VOLLETH & HELLER 1994; ROEHRS et al. 2010) justified the designation as separate „Hypsogine Group” subtribe (PAVLINOV & LISSOVSKY 2012). For many decades only the polytypic *Pipistrellus* (*Hypsugo*) *savii* (Bonaparte, 1837) with three subspecies, namely *P. s. savii, P. s. alaschanicus* (Bobrinskii, 1926) und *P. s. caucasicus* (Satunin, 1901), has been recognized for Europe and Central Asia (including Mongolia).

According to the current knowledge, the genus *Hypsugo* includes at least 17 species with Palearctic distribution (GRIMMBERGER & RUDLOFF 2009; 16 to 20 species according to Wikipedia 2020). The species and subspecies recognition within the genus *Hypsugo* with its morphologically difficult to distinguishable and cryptic forms as well as their distribution ranges are under discussion for more than 100 years (HORÁČEK et al. 2000; HORÁČEK 2004; GRIMMBERGER & RUDLOFF 2009; PAVLINOV & LISSOVSKY 2012). Despite the ongoing taxonomic debate, *H. alaschanicus* is now been widely accepted as a valid *Hypsugo* species (HORÁČEK & HANÁK 1986; NOWAK 1991; HORÁČEK et al. 2000; DOLCH et al. 2007; DATZMANN et al. 2012). Nonetheless and as result of the different taxonomic viewpoints, *H. alaschanicus* is often listed in older literature as *Vespertilio savii alaschanicus* or *Pipistrellus savii alaschanicus* with its probable synonym *Amblyotus velox* (CORBET 1978; KOOPMAN 1994; HORÁČEK et al. 2000).

Two *Hypsugo* species have been described for Mongolia, namely *H. alaschanicus* (SOKOLOV & ORLOV 1980; DOLCH et al. 2007; DATZMANN et al. 2012) und *H. savii* (STUBBE & CHOTOLCHU 1968; SOKOLOV & ORLOV 1980) with its subspecies *H. s. caucasicus*. SOKOLOV & ORLOV (1980) list a total of four records for both *Pipistrellus savii alaschanicus* und *P. s. caucasicus*, but does not assign the individual records to subspecies (nowadays species) level. Except for addressing the taxonomic reassignments within the *Vespertilioninae*, not much further progress towards unravelling the distribution and the status of *Hypsugo* species in Mongolia have been made for decades.

While both *Hypsugo* species are recognized for Mongolia, recent reports only describe new and further records of *H. alaschanicus* (DOLCH et al. 2007; DATZMANN et al. 2012). Within the years 2009 – 2011 a total of more than 50 individuals, primarily adult females and juveniles, have been captured at Šutegijn Bajan-gol in the south of Mongolia (Stubbe pers. comm). These records as well as captured juveniles in 2014 and 2017 describe the successful and continuous reproduction of *H. alaschanicus* in Mongolia.

Despite the increasing knowledge regarding the status of *H. alaschanicus*, a large uncertainty towards the status and distribution of *H. savii* (*H. s. caucasicus*) in Mongolia exists. With the increasing number of *Hypsugo* records the question arise, if *H. savii* has been simply overlooked or captured individuals were “wrongly” assigned to *H. savii* instead of *H. alaschanicus* or *vice versa* over the years.

The aim of this work is to gain a deeper understanding of the distribution and status of *Hypsugo* species in Mongolia. As a result, a new *Hypsugo* species named *H. stubbei* sp. nov. is described, which reveals large genetic distances to *H. savii* and *H. alaschanicus*.

## Materials and methods

Specimen data and sample material utilised in this study were collected over several decades in the frame of Mongolian-German Biological Expeditions since 1962 (e.g. STUBBE et al. 2005, 2012, 2017, 2020). Members of the Regional Committee of Mammalogy (LfA Säugetierkunde) Brandenburg-Berlin and Mongolian colleagues (e.g. DOLCH et al. 2007, 2021; SCHEFFLER et al. 2010, 2016) collected a large amount of data and material within the «Chiroptera Mongolia» field expeditions (since 1999). This resulted in a total of more than 1,800 records of bat specimens from more than 131 Mongolian capture locations (DOLCH et al. 2007, 2021) of which 83 specimens from 11 locations were assigned to the genus *Hypsugo* (fig. 1). The majority of the fieldwork was conducted throughout June (May) to September (October) within the respective years. Bats were captured primarily by mist-netting in potential hunting grounds along river or mountain valleys using specialized nets of various length and heights as previously described (DOLCH et al. 2007, 2021). All captured specimens were identified to species level in the field using available keys and descriptions (e.g. DOLCH et al. 2007). Measurements and morphological assessments were performed as previously described (DOLCH et al. 2007, 2021).

**Fig. 1:**
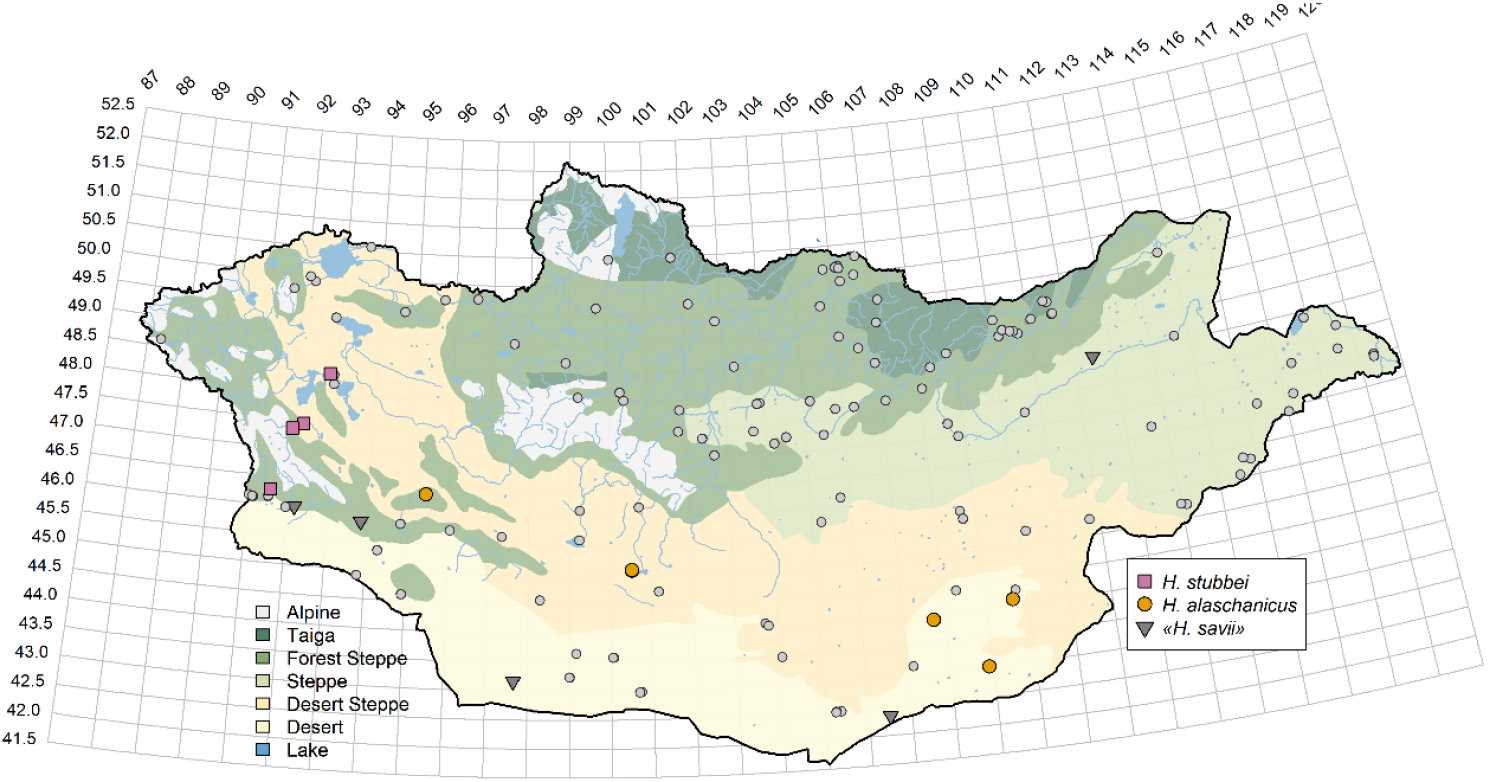
Records of *Hypsugo* species in Mongolia. The species are encoded by different colours and shapes as given in the right legend. The grey-coloured small circles depicting the collecting localities in Mongolia of the two main field expedition series (see material and methods). The grey-coloured triangles representing «*H. savii*» knowledge derived from literature (STUBBE & CHOTOLCHU 1968; SOKOLOV & ORLOV 1989; DAVAADORJ 2000 (in LKHAGVASUREN & SAMIYA 2004); TINNIN et al. 2002; see DOLCH et al. 2007) with approximate location assignment. The genus name of «*H. savii*» has been updated from *Pipistrellus savii* and/or *Vespertilio savii* in the source publications. The vegetation zones are colour-coded according to the left-side legend. The distribution map was created using R with modified shapefile data based on FREE (2018). Nearby locations may not be clearly visible as distinct points.

Samples for genetic analyses were taken from the wing membrane of capture living specimens using a sterile 3-mm diameter biopsy punch and stored in 70 % pure ethanol. Total DNA extraction from wing biopsy punches or chest muscle material (∼10 mg) from already collected museum specimens was performed using DNeasy Blood & Tissue Kit (Qiagen) according to the manufacturer protocol. DNA concentration in ng / μL was measured using a NanoDrop 2000 Spectrophotometer (Thermo Fisher Scientific). The primers for complete mitochondrial *NADH DEHYDROGENASE SUBUNIT1* (*mt-ND1*) gene amplification were ER66 and ER70 (MAYER & VON HELVERSEN 2001). For the partial mitochondrial *CYTOCHROME OXIDASE SUBUNITI* (*mt-COI*) amplification, a previously described vertebrate M13-tailed primer cocktail (C_VF1LFt1 + C_VR1LRt1) was used (IVANOVA et al. 2012). All amplifications were carried out in a total reaction volume of 25 µL containing 1.5 mM MgCl_2_ (*mt-COI*: 1.5-2 mM), 0.2 mM of each dNTP, 0.2 µM (*mt-COI*: 0.1-0.2 µM) of each primer, 0.75 units HGS Diamond Taq Polymerase and its reaction buffer including 2 µL (1-10 µL) DNA extract.

The PCR for both genes was performed separately using a Thermal Cycler (Analytik Jena) with an initial 10 min polymerase activation and DNA denaturation step at 95 °C and finished after 40 cycles by a final extension step at 72 °C for 10 min with a final hold at 4 °C. For *mt-ND1*, other cycling parameters were as described (MAYER & VON HELVERSEN 2001) with exception of the annealing (45 s) and elongation (105 s) time. For *mt-COI*, all other cycling parameters were in accordance with IVANOVA et al. (2012) but with an annealing time of 45 s. Amplified PCR products were checked using gel electrophoresis for size and quality. Upon success, PCR reactions were than purified using either ExoSAP-IT (Affymetrix, for *mt-ND1*) or NucleoSpin PCR Clean-up (Macherey-Nagel, for *mt-COI*) according to the respective manufacturer protocol. Forward and reverse sequencing was performed using standard Sanger sequencing method (MWG) with primers ER70 and ER89 for *mt-ND1* (MAYER & VON HELVERSEN 2001) and M13F and M13R for *mt-*COI (IVANOVA et al. 2012). Primer walking (5′-GGGCAGTAGCCCAAACTATTTC-3′) has been used to obtain the full *mt-ND1* sequence for *Hypsugo stubbei* sp. nov.. The lack of insertions, deletions and stop codons in the translated sequences support the mitochondrial origin of all obtained nucleotide sequences.

If not otherwise stated, all sequence analyses were performed in R (https://www.r-project.org/) with main algorithms from the following R packages: ‘phangorn’ (SCHLIEP 2011), ‘seqinr’ (CHARIF & LOBRY 2007), ‘ape’ (PARADIS & SCHLIEP 2019), ‘splits’ (EZARD et al. 2017), and ‘pegas’ (PARADIS 2010). Multiple sequence alignments were built using the web interface available at https://www.ebi.ac.uk/Tools/msa/clustalo/ (MADEIRA et al. 2019) of the Clustal Omega algorithm (SIEVERS et al. 2011; SIEVERS & HIGGINS 2018). The *mt-ND1* alignment was limited to the first 798 nucleotides from the ATG start codon. The *mt-COI* sequences and alignment were utilized for GenBank searches but not used further for in-depth phylogenetic analyses. Uncorrected genetic (*p*-) distances were calculated as percentages. Maximum-likelihood (ML), maximum-parsimony (MP), and neighbour-joining (NJ) algorithms with 1,000 bootstrap replications and Kimura-2-parameter (K2P, Kimura 1980) were used to reconstruct the phylogenetic evolution of *Hypsugo* species. Genetic clusters were delimited with the ‘Generalized Mixed Yule-Coalescent’ (GMYC) approach (PONS et al. 2006) as described by DATZMANN et al. (2012) using a single-threshold version.

Sequence data were submitted to NCBI GenBank under the accession numbers MW367760 – MW367764 (*mt-ND1*) and MW367765 - MW367769 (*mt-COI*). Third-party NCBI sequences used in this study are designated by their NCBI sequence number at appropriate places. *Miniopterus schreibersi* (KJ948260) and *Tadarida teniotis* (DQ915087) were used as outgroups.

All statistical analyses and visualizations were performed using R software (https://www.r-project.org/).

## Results and discussion

To gain a better understanding of the distribution of the genus *Hypsugo* in Mongolia, known species records that had already been flagged as somewhat “unusual” *H. alaschanicus* or yet undetermined *Hypsugo* species were re-evaluated within this study. Considering the likelihood of similarity in external characteristics, the main emphasis was put on records with either available genetic sample material or collected specimens. Of particular interest were 10 specimens captured at three different locations during the 2011 «Chiroptera Mongolia» field expedition (fig. 1, table 1). These specimens were initially characterized as somewhat bright coloured *Hypsugo* sp. and provisionally assigned to *H. alaschanicus* taking into account the colour-variation and two known colour variants already described for Mongolia (DOLCH et al. 2007). In total eight adult male were captured swarming around a potential hibernating roost as well as one adult and one juvenile female at two further locations (see table 1). The adult female clearly showed signs of having an offspring in the current year. A specimen (voucher Stubbe 2002/181) from 2002 was additionally included (table 1), that was captured in the same area were the second record of (at that time) *P. savii caucasicus* for Mongolia was detected in 1964 (STUBBE & CHOTOLCHU 1968).

**Table 1a:** Species records of *Hypsugo stubbei* sp. nov..

| Specimen | Sex | Age | Collector | Date | Location | GEO Reference | Reproduction | Forearm [mm] |
| --- | --- | --- | --- | --- | --- | --- | --- | --- |
| 805 | ♀ | ad. | Chiroptera Mongolia | 17.07.2011 | Tsenkher-gol | 47°26'37.7"N / 92°13'45.8"E | + | 34.5 |
| 838 | ♂ | ad. | Chiroptera Mongolia | 18.07.2011 | Khoya Tsenkheiyin Aguy | 47°20'55.7"N / 91°57'04.6"E | + | 34.8 |
| 839 | ♂ | ad. | Chiroptera Mongolia | 18.07.2011 | " | " | + | 33.8 |
| 840 | ♂ | ad. | Chiroptera Mongolia | 18.07.2011 | " | " | + | 34.1 |
| 824 | ♂ | ad. | Chiroptera Mongolia | 18.07.2011 | " | " | - | 34.8 |
| 825 | ♂ | ad. | Chiroptera Mongolia | 18.07.2011 | " | " | + | 34.9 |
| 854 | ♂ | ad. | Chiroptera Mongolia | 18.07.2011 | " | " |  | 33.0 |
| 855 | ♂ | ad. | Chiroptera Mongolia | 18.07.2011 | " | " |  | 35.0 |
| 856 | ♂ | ad. | Chiroptera Mongolia | 18.07.2011 | " | " |  | 33.1 |
| 858 | ♀ | juv. | Chiroptera Mongolia | 22.07.2011 | Gozgor Ulaan-uul | 46°16'06.4"N / 91°32'44.7"E |  |  |
| 1012 | ♀ | ad. | Stubbe | 25.08.2002 | Čonocharajchijn-gol | 48°20'N / 92°48'E |  | 36.0 |

**Table 1b:** Species records of *Hypsugo alaschanicus* used in genetic analysis.

| Specimen | Sex | Age | Collector | Date | Location | GEO Reference | Reproduction | Forearm [mm] |
| --- | --- | --- | --- | --- | --- | --- | --- | --- |
| 312 | ♂ | ad. | Chiroptera Mongolia | 26.08.2005 | Orog Nuur | 45°04'45.5"N / 100°32'07.6"E | + | 33.3 |
| 947 | ♀ | juv. | Chiroptera Mongolia | 30.08.2014 | Hatni Bulag | 43°00'13.3"N / 108°54'31.7"E |  | 35.3 |
| 1005 | ♀ | ad. | Stubbe | 03.08.2009 | Šutejin Bajan-gol | 43°54'19.3"N / 107°43'45.5"E |  | 35.0 |
| 1006 | ♀ | ad. | Stubbe | 24.07.2011 | " | " |  | 37.0 |
| 1007 | ♀ | ad. | Stubbe | 13.07.2010 | " | " |  | 36.0 |
| 1013 | ♀ | ad. | Stubbe | 16.07.2007 | Chatanbulag | 43°54'20.0"N / 107°43'38.4"E | + | 36.0 |

In total 12 samples from six “unusual” and six typical *H. alaschanicus* specimens captured at eight different locations throughout Mongolia were analysed genetically based on either the mitochondrial *NADH DEHYDROGENASE SUBUNIT1* (*mt-ND1*) or the mitochondrial *CYTOCHROME OXIDASE SUBUNITI* (*mt-COI*) gene sequences (table 2). Initial nucleotide (nt) BLAST searches against the NCBI GenBank database (last search 11.12.2020) with the “unusual” *H. alaschanicus* revealed substantial sequence differences to both known *H. savii* (1^st^ best hits) and *H. alaschanicus* (2^nd^ best hits) sequences. For the complete 957 nt of the *mt-ND1* gene the percent identity was about 91 % to *H. savii* and about 89 % to *H. alaschanicus*. The percent identity for the partial 657 nt of the *mt-COI* gene was about 92 % to *H. savii* and about 90 % to *H. alaschanicus*, respectively. These initial results already indicate a potential new, yet undiscovered cryptic *Hypsugo* species from Western Mongolia.

**Table 2:** List of *Hypsugo* specimens sequenced within this study. The specimens with the grey-coloured bold haplotype were used for phylogenetic analyses as representative case for each haplogroup. NCBI GenBank sequence submissions were only made for sequences obtained from either collected specimens (vouchers) or for a selected specimen of each haplogroup and combination.

| Specimen | Species | Sample | Haplotype Group |  | Geo Reference |  | NCBI GenBank |  | Voucher |
| --- | --- | --- | --- | --- | --- | --- | --- | --- | --- |
|  |  |  | *mt-ND1* | *mt-COI* | North | East | *mt-ND1* | *mt-COI* |  |
| **805** | *H. stubbei* sp. nov. | 076/11 | Hssp-HG01 | Hssp-HG01 | 47°26'37.7" | 92°13'45.8" |  |  |  |
| **824** | *H. stubbei* sp. nov. | 106/11 | Hssp-HG01 | Hssp-HG01 | 47°20'55.7" | 91°57'04.6" |  |  |  |
| **838** | *H. stubbei* sp. nov. | 103/11 | Hssp-HG01 | Hssp-HG01 | 47°20'55.7" | 91°57'04.6" |  |  |  |
| **840** | *H. stubbei* sp. nov. | 105/11 | Hssp-HG01 | Hssp-HG01 | 47°20'55.7" | 91°57'04.6" |  |  |  |
| **858** | *H. stubbei* sp. nov. | 141/11 | Hssp-HG01 |  | 46°16'06.4" | 91°32'44.7" |  |  |  |
| **1012** | *H. stubbei* sp. nov. | 181/2002 | **Hssp-HG01** | **Hssp-HG01** | 48°20' | 92°48' | MW367764 | MW367769 | 2002/181 |
| **312** | *H. alaschanicus* | 26/05 | **Hala-HG03** | **Hala-HG01** | 45°04'45.5" | 100°32'07.6" | MW367760 | MW367765 | M.11252 |
| **947** | *H. alaschanicus* | 094/14 | **Hala-HG02** | **Hala-HG02** | 43°00'13.3" | 108°54'31.7" | MW367761 | MW367766 | M.11324 |
| **1005** | *H. alaschanicus* | 29/2009 | **Hala-HG04** | Hala-HG01 | 43°54'19.3" | 107°43'45.5" | MW367762 | MW367767 |  |
| **1006** | *H. alaschanicus* | 14/2011 | Hala-HG04 | Hala-HG01 | 43°54'19.3" | 107°43'45.5" |  |  |  |
| **1007** | *H. alaschanicus* | 9/2010 | Hala-HG04 | Hala-HG01 | 43°54'19.3" | 107°43'45.5" |  |  |  |
| **1013** | *H. alaschanicus* |  | **Hala-HG01** | Hala-HG01 | 43°54'20.0" | 107°43'38.4" | MW367763 | MW367768 |  |

Phylogenetic trees using different algorithms and K2P distances (fig. 2, see material and methods section) were re-constructed to deepen the insights into the phylogenetic relationship of Mongolian *Hypsugo* species. These analyses revealed that the Mongolian bats of the genus *Hypsugo* fall into two largely different clades. One well-supported clade represent *H. alaschanicus* which had a *mt-ND1* sequence divergence of up to 0.2% and bootstrap supports of 90 to 100 in the various reconstruction approaches (fig. 2, table 3). The most closely related Western Palaearctic species differed by at least 9 % *mt-ND1* sequence divergence to both, *H. savii* and *H*. cf. *darwinii*, respectively (table 3). This is in well agreement with previous phylogenetic reconstructions of *H. alaschanicus* sequences from Mongolia (Datzmann et al. 2012). A second clade represents *Hypsugo* sequences from Western Mongolia which formed a single haplotype with all sequences being identical. The most closely related *Hypsugo* species differs by at least 8.4 % for *H*. cf. *darwinii* followed by 9 % for *H. savii mt-ND1* sequence divergence (table 3). The difference between the two Mongolian species is at least 11.3 % *mt-ND1* sequence divergence (table 3). Species delineation using GMYC method also uncovered the two clusters (fig. 2). Statistical parsimony minimum-spanning tree reconstructing using *p*-distances on the closely related Western Palearctic species revealed also substantial genetic differences of the Western Mongolian *Hypsugo* samples to *H. alaschanicus* and to *H. savii* (fig. 3). In the latter analysis, 41 theoretical haplotypes are missing to the closest, and in this case Western Palearctic, species *H. savii*. The closest clade comprises only *H. savii* sequence derived from the middle east, namely sequence records from Israel, Turkey, and Iran. As these origins already overlap with described distribution range of *H. s. caucasicus* (DIETZ & KIEFER 2014), it is possible that this clade may represent *H. s. caucasicus* but an alternative *H. savii* subspecies cannot be unambiguously excluded.

**Table 3:** *mt-ND1* inter- and intraspecific genetic (*p*-) distances of the genus *Hypsugo*. The intraspecific distances are given in bold at the grey-coloured diagonal as mean and standard deviation. The lower triangle contains the average and standard deviations between each species or subspecies while the upper part indicating the minimum – maximum range. All distances are uncorrected pairwise *p*-distances in percentage. Missing estimates are indicated by n/a. The number of analysed unique sequences (n) are given. The assignment of subspecies names to the individual *H. savii* clades and geographic groups must be taken with caution and are assigned here to aid interpretation. The upper table contains the distances based on currently designated species level while the lower table expands towards genetic *H. savii* clades, aka potential subspecies.

| Lineage | Abbr. | n | Hstu | Hala | Hsav | Hdar | Hari | Hcad | Hsp1 |
| --- | --- | --- | --- | --- | --- | --- | --- | --- | --- |
| *H. stubbei* sp. nov. | Hstu | 1 | n/a | 11.3 - 11.8 | 9.0 - 11.5 | 8.4 - 9.7 | 12.4 - 12.5 | 14.0 - 14.0 | 17.4 - 17.4 |
| *H. alaschanicus* | Hala | 6 | 11.4 ± 0.2 | **0.2 ± 0.1** | 9.0 - 11.8 | 9.0 - 10.4 | 13.6 - 14.4 | 14.9 - 15.2 | 10.7 - 16.7 |
| *H. savii* | Hsav | 10 | 10.0 ± 0.8 | 10.1 ± 0.6 | **4.2 ± 3.1** | 7.6 - 11.4 | 10.2 - 13.1 | 11.9 - 13.0 | 13.9 - 16.3 |
| *H. cf. darwinii* | Hdar | 9 | 9.2 ± 0.4 | 9.7 ± 0.3 | 9.1 ± 0.7 | **1.2 ± 0.5** | 12.8 - 13.9 | 12.5 - 13.4 | 16.7 - 17.8 |
| *H. ariel* | Hari | 2 | 12.4 ± 0.0 | 13.9 ± 0.3 | 12.0 ± 0.9 | 13.4 ± 0.3 | **0.4 ± 0.0** | 14.8 - 15.0 | 15.3 - 15.7 |
| *H. cadornae* | Hcad | 1 | 14 | 15.1 ± 0.1 | 12.3 ± 0.3 | 12.8 ± 0.3 | 14.9 ± 0.1 | n/a | 15.9 - 15.9 |
| *H. sp.* | Hsp1 | 1 | 17.4 | 15.6 ± 2.4 | 15.2 ± 0.8 | 17.1 ± 0.5 | 15.5 ± 0.3 | 15.9 | n/a |

| Lineage | Abbr. | n | Hstu | Hala | Hsa1 | Hsa2 | Hsa3 | Hdar | Hari | Hcad | Hsp1 |
| --- | --- | --- | --- | --- | --- | --- | --- | --- | --- | --- | --- |
| *H. stubbei* sp. nov. | Hstu | 1 | n/a | 11.3 - 11.8 | 10.2 - 10.8 | 9.5 - 11.5 | 9.0 - 9.6 | 8.4 - 9.7 | 12.4 - 12.5 | 14.0 - 14.0 | 17.4 - 17.4 |
| *H. alaschanicus* | Hala | 6 | 11.4 ± 0.2 | **0.2 ± 0.1** | 9.3 - 10.8 | 9.0 - 11.8 | 9.5 - 10.6 | 9.0 - 10.4 | 13.6 - 14.4 | 14.9 - 15.2 | 10.7 - 16.7 |
| *H. savii savii* | Hsa1 | 3 | 10.5 ± 0.3 | 10.3 ± 0.4 | **0.4 ± 0.2** | 1.8 - 3.9 | 6.0 - 7.6 | 7.6 - 11.3 | 12.5 - 13.1 | 11.9 - 12.1 | 16.0 - 16.3 |
| *H. savii caucasicus* (?) | Hsa2 | 4 | 10.1 ± 1.0 | 10.0 ± 0.8 | 2.4 ± 0.6 | **0.6 ± 0.2** | 6.0 - 10.4 | 8.0 - 11.4 | 11.8 - 12.6 | 12.1 - 12.6 | 13.9 - 15.5 |
| *H. savii ochromixtus* | Hsa3 | 3 | 9.3 ± 0.3 | 10.0 ± 0.3 | 6.9 ± 0.5 | 7.6 ± 1.4 | **0.5 ± 0.1** | 8.4 - 9.4 | 10.2 - 11.3 | 12.2 - 13.0 | 15.3 - 15.3 |
| *H. cf. darwinii* | Hdar | 9 | 9.2 ± 0.4 | 9.7 ± 0.3 | 9.3 ± 0.8 | 9.2 ± 0.9 | 8.9 ± 0.3 | **1.2 ± 0.5** | 12.8 - 13.9 | 12.5 - 13.4 | 16.7 - 17.8 |
| *H. ariel* | Hari | 2 | 12.4 ± 0.0 | 13.9 ± 0.3 | 12.8 ± 0.2 | 12.3 ± 0.3 | 10.9 ± 0.4 | 13.4 ± 0.3 | **0.4 ± 0.0** | 14.8 - 15.0 | 15.3 - 15.7 |
| *H. cadornae* | Hcad | 1 | 14 | 15.1 ± 0.1 | 12.0 ± 0.1 | 12.3 ± 0.2 | 12.5 ± 0.4 | 12.8 ± 0.3 | 14.9 ± 0.1 | n/a | 15.9 - 15.9 |
| *H. sp.* | Hsp1 | 1 | 17.4 | 15.6 ± 2.4 | 16.1 ± 0.2 | 14.6 ± 0.7 | 15.3 ± 0.0 | 17.1 ± 0.5 | 15.5 ± 0.3 | 15.9 | n/a |

**Fig. 2:**
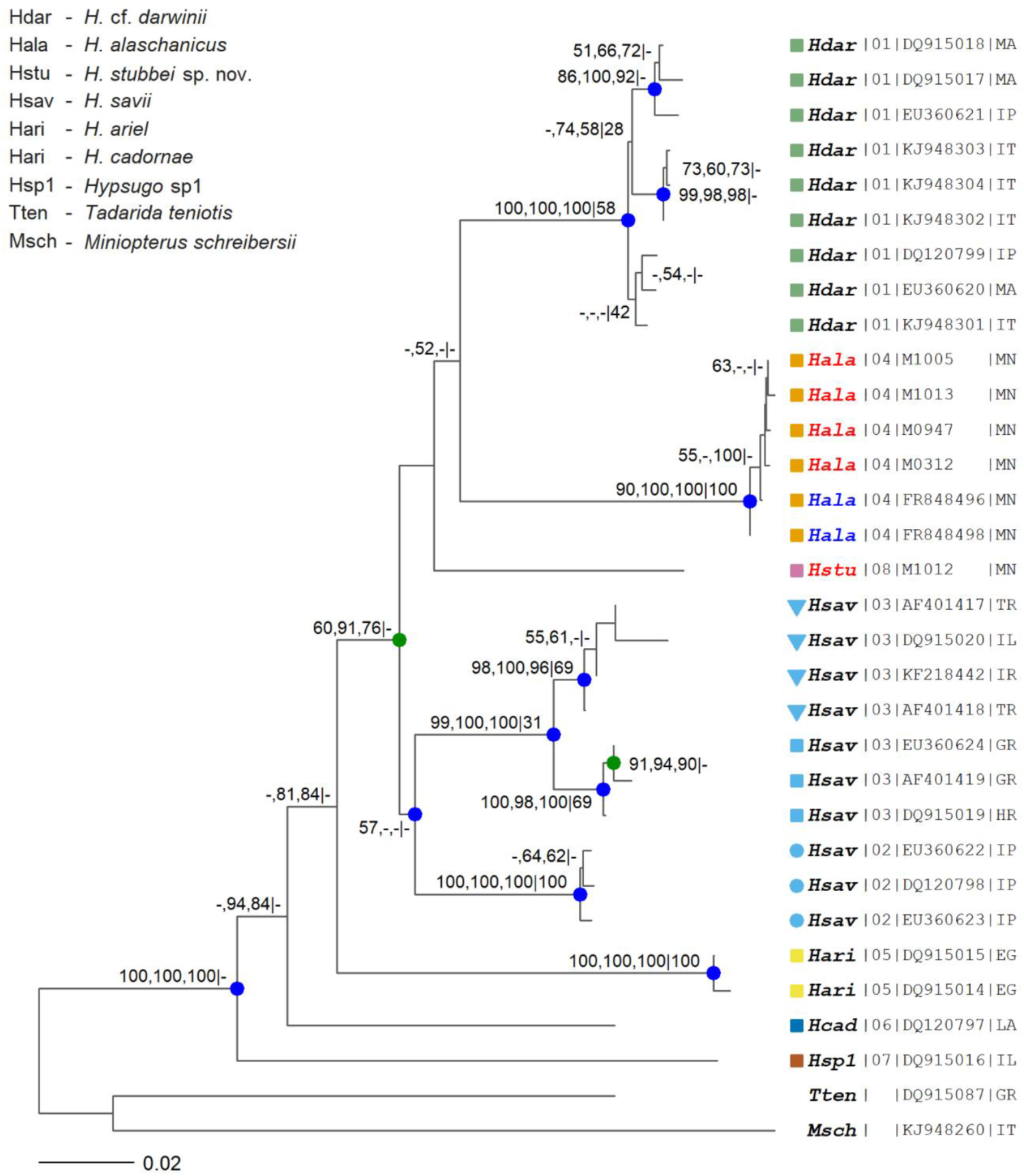
Neighbour-joining (NJ) tree based on the alignment of the first 798 nucleotides of the mitochondrial *ND1* gene (subunit one of NADH dehydrogenase) estimated using Kimura’s 2-Parameter (K2P) distances. The values on the nodes and branching points are bootstrap support values (>50%) of the various algorithms (NJ, maximum-likelihood (ML), maximum-parsimony (MP)) followed by the GMYC / Yule support (0-100) for species delineation. The leaf nodes tips are labelled based on the species assignment using different colours and in case of potential subspecies different shapes with identical colours. The 4-letter species abbreviations (see inset legend for meaning) in red (this study) and blue (published NCBI GenBank sequences) colour represent sequences from specimen material collected in Mongolia (MN). The species abbreviations are followed by the GMYC / Yule cluster number, the NCBI GenBank number (in case of published sequences) or the specimen number (M) and finally the 2-letter country code. Branching points with statistical relevant GMYC / Yule posterior probabilities are marked by color-coded circles as follows: orange ≥ 0.9, green ≥ 0.95, blue ≥ 0.99. *M. schreibersii* and *T. teniotis* functioning as outgroup species.

**Fig. 3:**
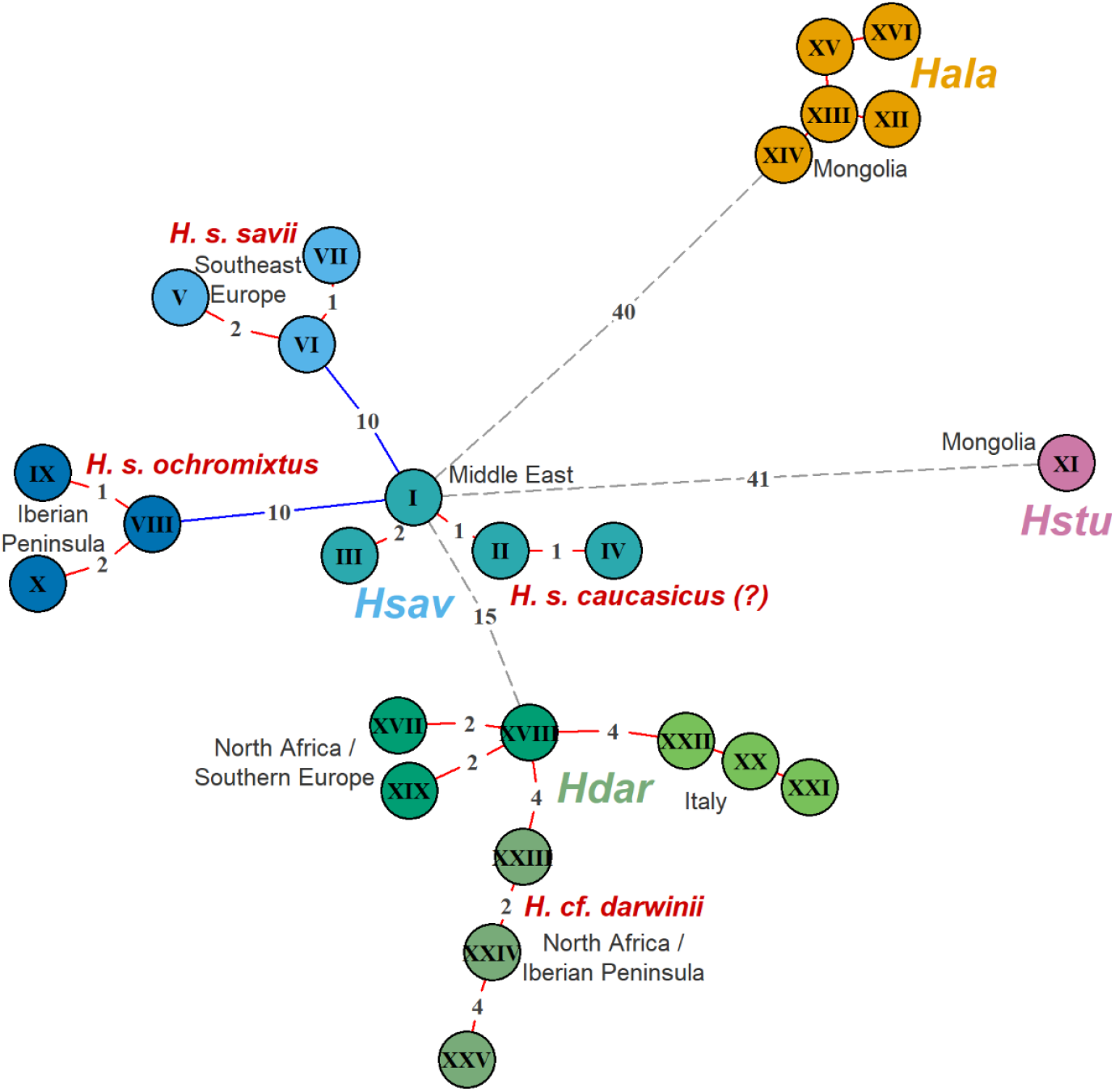
Statistical parsimony minimum-spanning tree (MST) based on (uncorrected) *p*-distances estimated from the alignment of the first 798 nucleotides of the mitochondrial *ND1* gene of selected *Hypsugo* species with known or potential distribution in Mongolia. Theoretical missing haplotypes are indicated by numbers in the connecting line of the minimum parsimonious path. The coloured circles indicating the different species (and potential subspecies or genetic clades) as follows: *H. stubbei* sp. nov. (Hstu) = magenta, *H. alaschanicus* (Hala) = yellow, *H. savii* (Hsav) = blue / blue-green, and *H*. cf. *darwinii* = greenish. The line colour of the connecting lines reflecting the significance level (*p*) of connectivity between genetically similar haplotypes with red (*p* < 0.01), blue (*p* < 0.05) and grey – not significant (*p* ≥ 0.05). The assignment of subspecies names to the individual *H. savii* clades and geographic groups must be taken with caution and are assigned here to aid interpretation. The assignment of *H. s. caucasicus* is based on the overlap of the described distribution range and the capture location of the haplotype under investigation.

Nonetheless, the substantial sequence divergence and statistical support of the phylogenetic reconstruction clearly shows the existence of a novel yet undiscovered and unrecognized cryptic *Hypsugo* species from Western Mongolia. To indicate this, we designate these specimens as *H. stubbei* sp. nov. named after Prof. Dr. Michael Stubbe. He has made significant contributions into the biological exploration of Mongolia for over 60 years. With the discovery of *Pipistrellus* (today *Hypsugo*) in 1964 in Mongolia (STUBBE & CHOTOLCHU 1968), he pinpointed on recognizable difference in the hairiness of the tail requiring further investigations for unambiguous species delineation. All these suggestions have contributed to the discovery of this cryptic species after more than 50 years.

### Hypsugo stubbei sp. nov

#### Holotyp

2002/181; Collection Stubbe; female adult; 25.08.2002; Čonocharajchijn-gol, N048°20’, E092°48’; Mongolia; Skull, Skin and Body (Specimen 1012, see fig. 4).

**Fig. 4:**
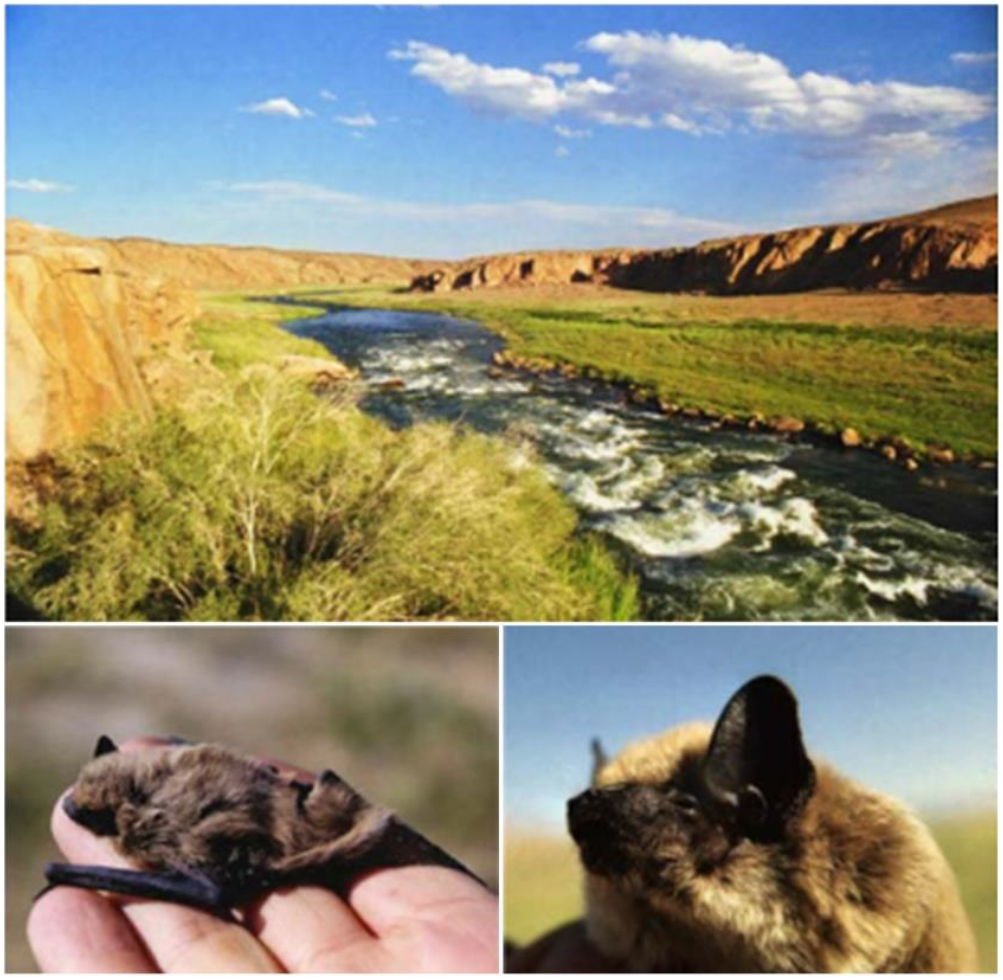
*Hypsugo stubbei* sp. nov. type locality from Čonocharajchijn-gol. Typus (2002/181; specimen 1012) and *Terra typica* at Čonocharajchijn-gol 2002, a capture locality of *Hypsugo stubbei* sp. nov., *Myotis davidii, Myotis petax, Eptesicus gobiensis* and *Plecotus kozlovi*. Photos: M. STUBBE.

#### Measures

The measurements of the holotype are as follows: mass 7 g, forearm 36 mm, head-body-length 55 mm, tail length 40 mm, tibia length 34 mm, ear length 13 mm, and tragus length 5.5 mm. The skull measurements are: condylobasal length 13.3 mm, zygomatic width 8.8 mm, interorbital constriction width 3.6 mm, UK (Length of mandible from symphysis to condylar process) 9.9 mm, CORH (UA, Hight of coronoid process) 3.1 mm, Length of upper tooth row from canine to third molar 4.5 mm, and Length of lower tooth row from canine to third molar 4.9 mm. Despite large genetic differences to *H. alaschanicus* and *H. savii* no substantial differences between the species and subspecies were found regarding skull as well as common external measures (fig. 5, 6).

**Fig. 5:**
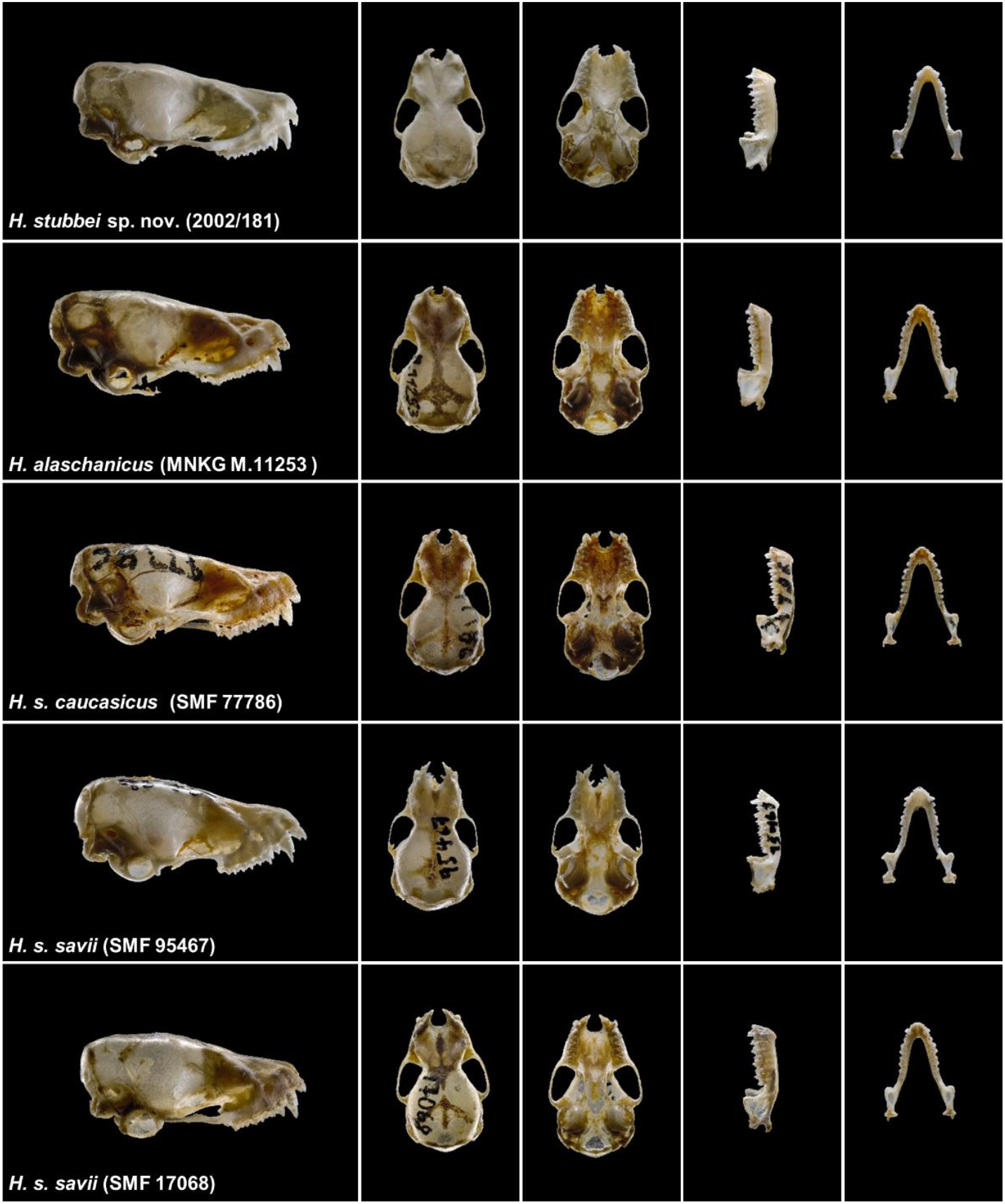
Skulls of selected *Hypsugo* species and subspecies. From top to bottom: *H. stubbei* sp. nov (Collection Stubbe 2002/181, Specimen 1012, Mongolia), *H. alaschanicus* (Senckenberg Museum of Natural History Goerlitz, Specimen 313, Mongolia), *H. savii caucasicus* (Specimen 1584, Kyrgyzstan) und *H. savii savii* from Germany (Specimen 1629, Bavaria) and Italy (Specimen 1625, Sicily) (all Senckenberg Museum Frankfurt / Main). From left to right: The cranium in lateral, dorsal, and ventral view followed by the mandible in lateral and dorsal view. Photos: B. GÄRTNER.

**Fig. 6:**
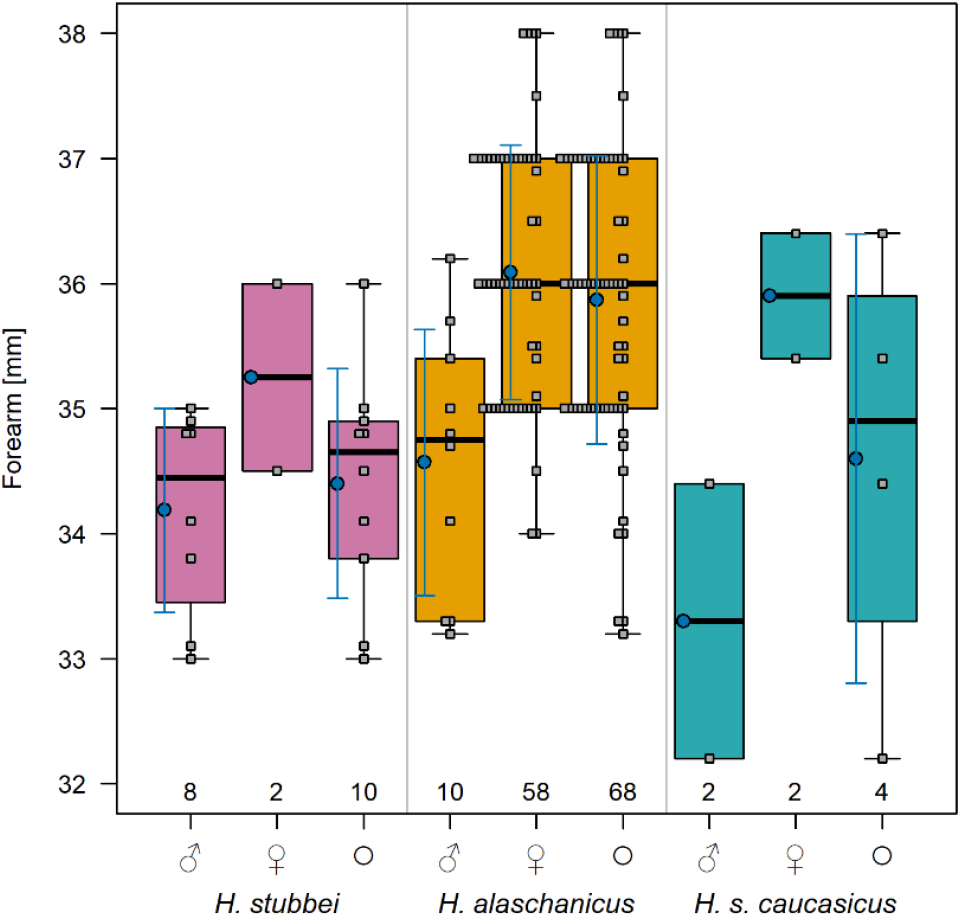
Boxplots and density plots of forearm size for species or subspecies of the genus *Hypsugo* from Mongolia (*H. stubbei* sp. nov, *H. alaschanicus*) or Kyrgyzstan (*H. savii caucasicus*). The measurements are separated according to the gender or total (+ *indet*) with the number of adult individuals (n) marked at the bottom. The mean (blue circle) and standard deviation (blue solid lines) are additionally depicted in each boxplot for n > 2. The grey-coloured squares indicating individual measurements.

#### Description

The initial designation of captured *H. stubbei* specimens as, even with doubts, “unusual” *H. alaschanicus* illustrate that morphological differences of this species to *H. alaschanicus* and *H. savii* are rather marginal or less obvious pronounced and thus more difficult to detect (fig. 5, 6; see table 4). However SOKOLOV & ORLOV (1980), which likely analysed *Hypsugo* specimens also from Mongolia, described for *H. alaschanicus* (named *Pipistrellus savii alaschanicus*) a brown-yellowish coloration of the dorsal fur with a brownish-yellowish underside. In contrast, specimens designated by them as *H. savii caucasicus* (named *P. s. caucasicus*) showed an overall more greyish coloration with a grey-straw yellow dorsal and a white ventral fur.

We also noticed differences in coloration as already mentioned above, which essentially confirm the descriptions by SOKOLOV & ORLOV (1980). *Hypsugo stubbei* is, as *H. alaschanicus*, a small-sized (see measurements above) contrast-rich species with a relatively long and dense pelage. The dorsal fur is two coloured with an almost black basis and a bright ochre upper part and hair tips showing a reddish tinge (fig. 4, 7, 8). *Hypsugo alaschanicus* in contrast, with a browner and a rather grey colour variant (DOLCH et al. 2007), shows a dorsal pelage with a black-brown hair basis and dark-brown to grey-brown upper part with light, white-yellowish hair tips (fig. 7, 9). The ventral fur of *H. stubbei* is clearly brighter with a light-grey whitish to white colour and darker hair basis (fig. 4, 7, 8). The transition into the slightly brighter ventral fur for *H. alaschanicus* is, in slight contrast, rather smooth with a light-grey to yellowish-grey hair colour with a grey-brown hair basis. Small differences are also observable regarding the coloration of the ears and wing membranes. The skin colour of the yet-captured *H. stubbei* specimens are dark-brown to almost black, while *H. alaschanicus* specimens from Mongolia show predominantly a light to dark brown, rarely an almost black, skin coloration. Together with the bright hair coloration, *H. stubbei* appears to be even more contrasty than the already contrast-rich *H. alaschanicus* (fig. 7 - 9). To what extend the differences in coloration are characteristic enough to distinguishing *H. stubbei* and *H. alaschanicus* must be further investigated as *H. alaschanicus* comprises broad colour variations with two observed colour variants (DOLCH et al. 2007, fig. 9).

**Fig. 7:**
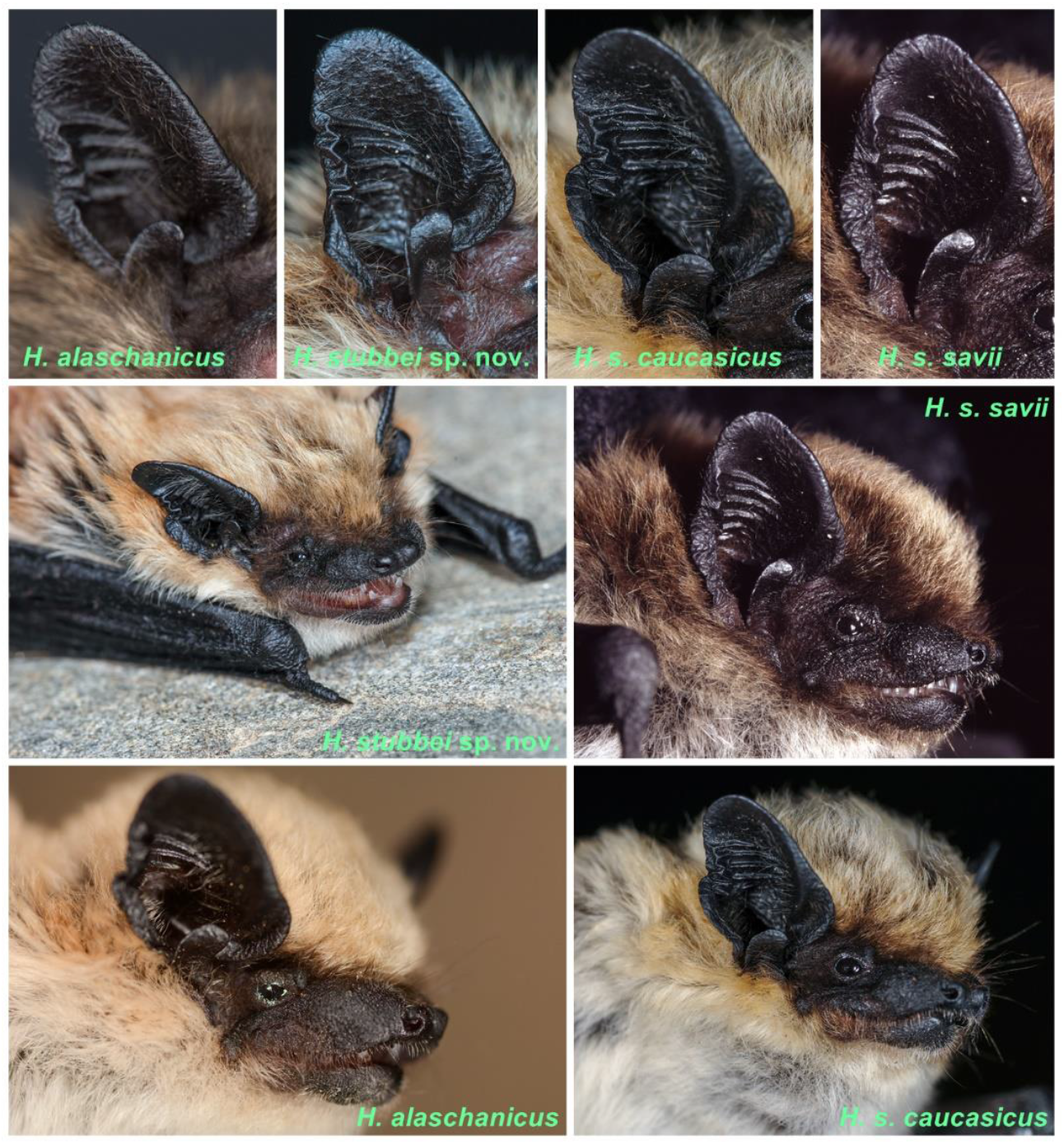
Ears and portraits of the four *Hypsugo* species or subspecies with known or potential distribution in Mongolia. The species or subspecies names are given as inset text. Photos: *H. stubbei* sp. nov. – D. STEINHAUSER; *H. alaschanicus* (ear) – B. GÄRTNER, *H. alaschanicus* (portrait), *H. savii savii* and *H. s. caucasicus* – C. DIETZ.

**Fig. 8:**
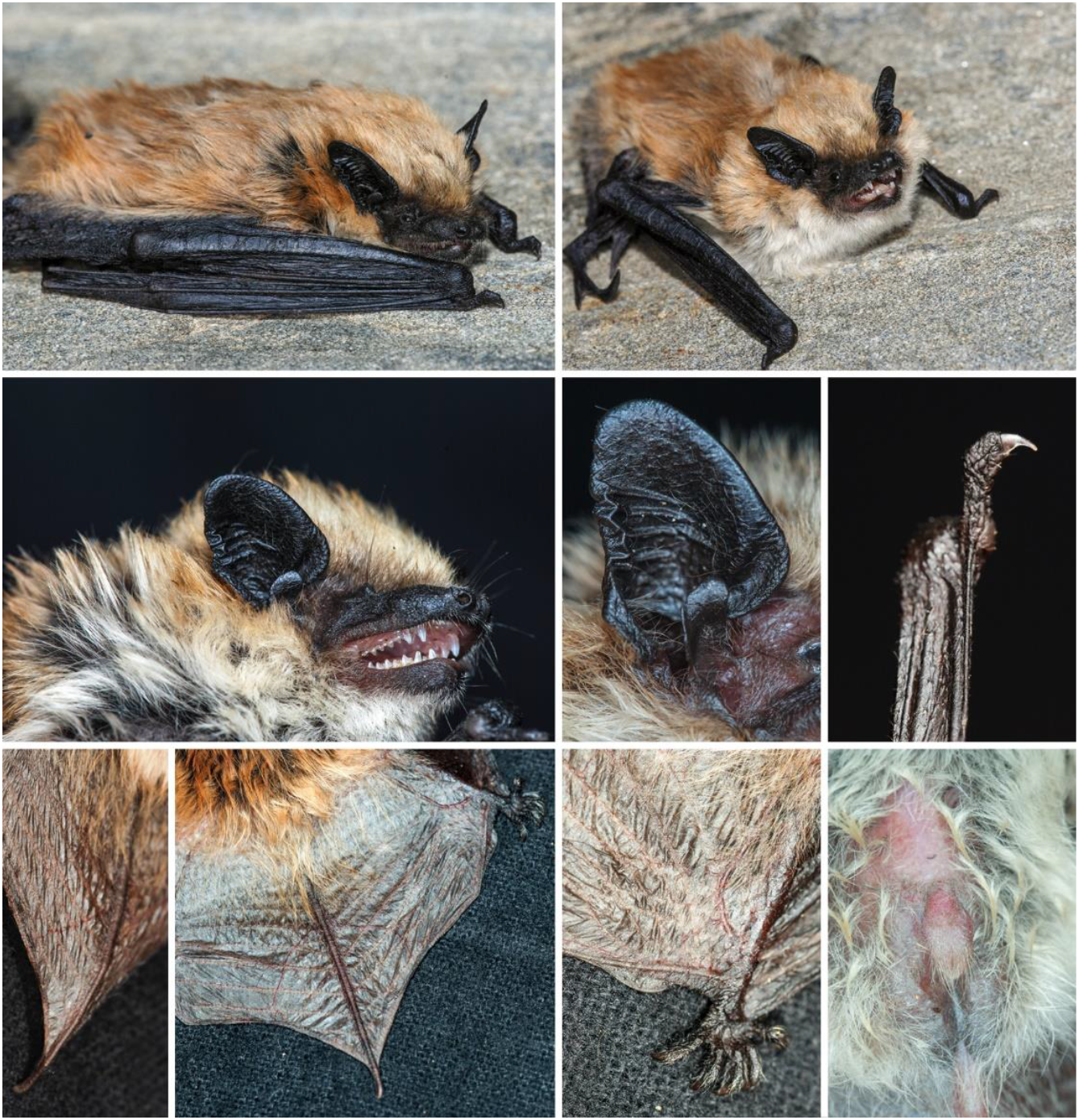
Overview pictures of *H. stubbei* sp. nov.. The full body pictures (top) clearly depict the colouration of the dorsal and ventral fur as well as the skin. The middle pictures show the portrait, the ear and claw. The bottom pictures depict the hair coverage of the tail, the epiblema and the penis. Ear and left tail image - Specimen 805, all other - Specimen 838. Photos: D. STEINHAUSER.

**Fig. 9:**
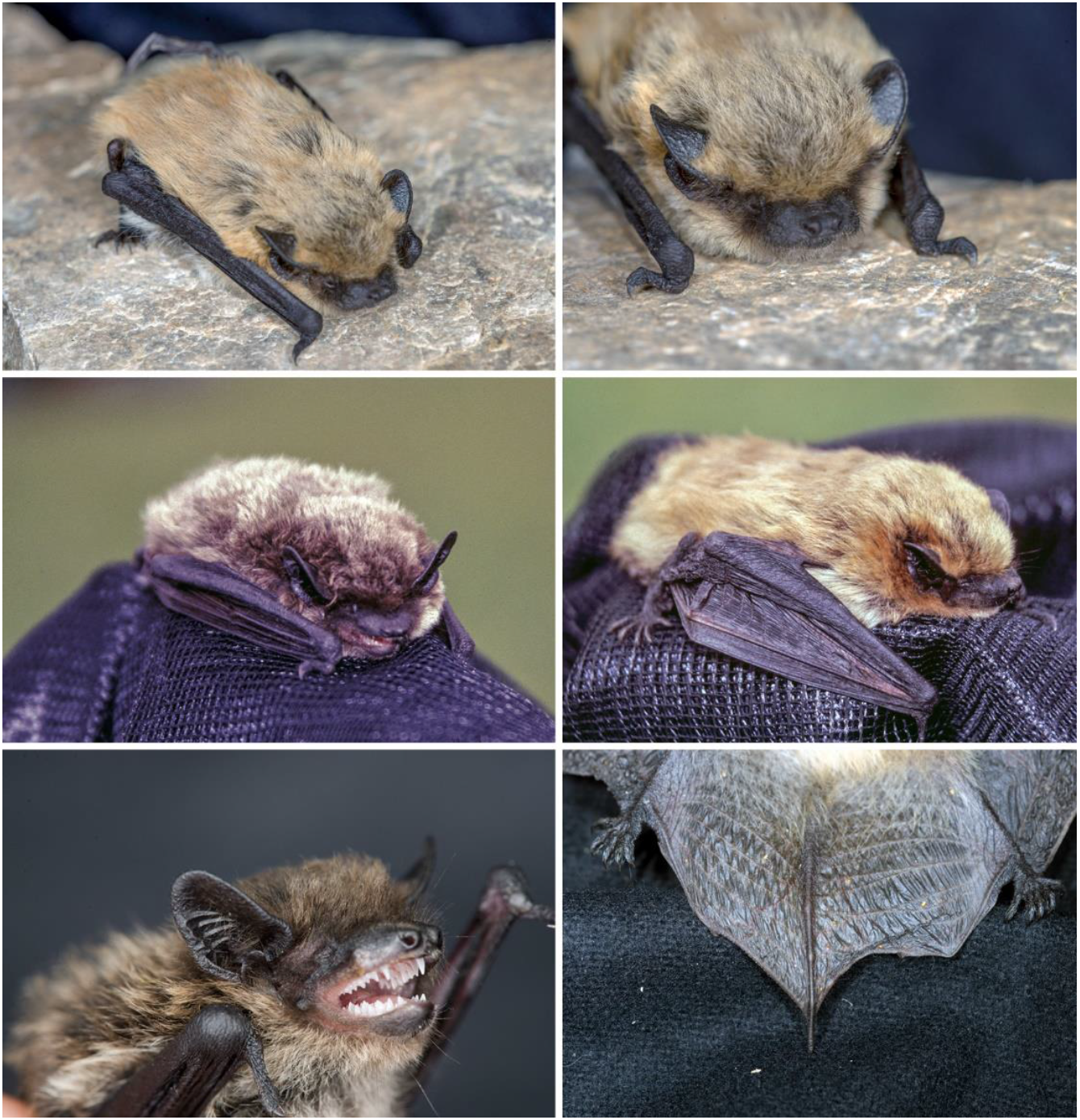
Overview pictures of *H. alaschanicus*. All pictures were taken at Orog-nuur in 2019 (top, bottom right) and 2005 (middle). The middle pictures show the two-colour variant of *H. alaschanicus* (see DOLCH et al., 2007). The bottom left pictures illustrates as browner colour variant from 2014. Photos: 2005 - D. STEINHAUSER, 2014 and 2019 - B. GÄRTNER.

STUBBE & CHOTOLCHU (1968) described that their captured *Hypsugo* specimens (designated as *P. s. caucasicus*) from Western Mongolia had a hairy upper part of the tail wing. This is in agreement with the captured and genetically analysed *H. stubbei* specimens from 2006 and 2011, which also have a narrow area of hairy skin (fig. 8, table 1). *Hypsugo* specimens from Kyrgyzstan that are designated as *H. savii caucasicus* (Senckenberg Museum Frankfurt) revealed in contrast to *H. stubbei* a much more hairy tail membrane. The hair coat reaches the heel, extends further along the tail spine and reveal hairy tibia. This has not been observed for the analysed *H. stubbei* specimens.

The hair tips of *H. s. caucasicus* are silver-white tinted according to DIETZ & KIEFER (2014) and thereby represent a good distinguishable characteristic to *H. s. savii* (fig. 7). As described above all *Hypsugo* specimens from Western Mongolia show a bright ochre to straw yellow dorsal fur coloration with a partial reddish tinge (fig. 7, 8) but we never observed a silver-white coloration.

#### Distribution

Currently, *H. stubbei* is known from four different locations within Mongolia with a total of 11 specimens including six specimens with genetic confirmation from all the four locations (fig. 1; table 1, 2). This species has currently only been caught in the far west of Mongolia covering the area near Bulgan Gol and the Altai Mountains (fig. 1). An overlap of the distribution area with *H. alaschanicus*, which in Mongolia is likely distributed more central south based on the few records (fig. 1), cannot currently be ruled out. The habitat of *H. stubbei* includes semi-deserts and steppes up to high mountain areas (fig. 10). For *H. savii* a very large distribution area is currently assumed with *H. s. caucasicus* apparently being mainly distributed throughout Central Asia, Iran, Turkmenistan, Afghanistan, Uzbekistan and Kyrgyzstan (DIETZ & KIEFER 2014). According to GVOZDEV & STRAUTMAN (1985) *H. s. caucasicus* is the only known *Hypsugo* subspecies from Kazakhstan. Based on their description the *Hypsugo* specimens reveal a brownish-ochre coloration of the dorsal fur apparently similar to the colorations of the one we found in western Mongolia. It could be possible, that the *Hypsugo* specimens of Kazakhstan are *H. stubbei*. As phylogenetic analyses from those specimens are according to our knowledge currently missing, we cannot further delineate this.

**Fig. 10:**
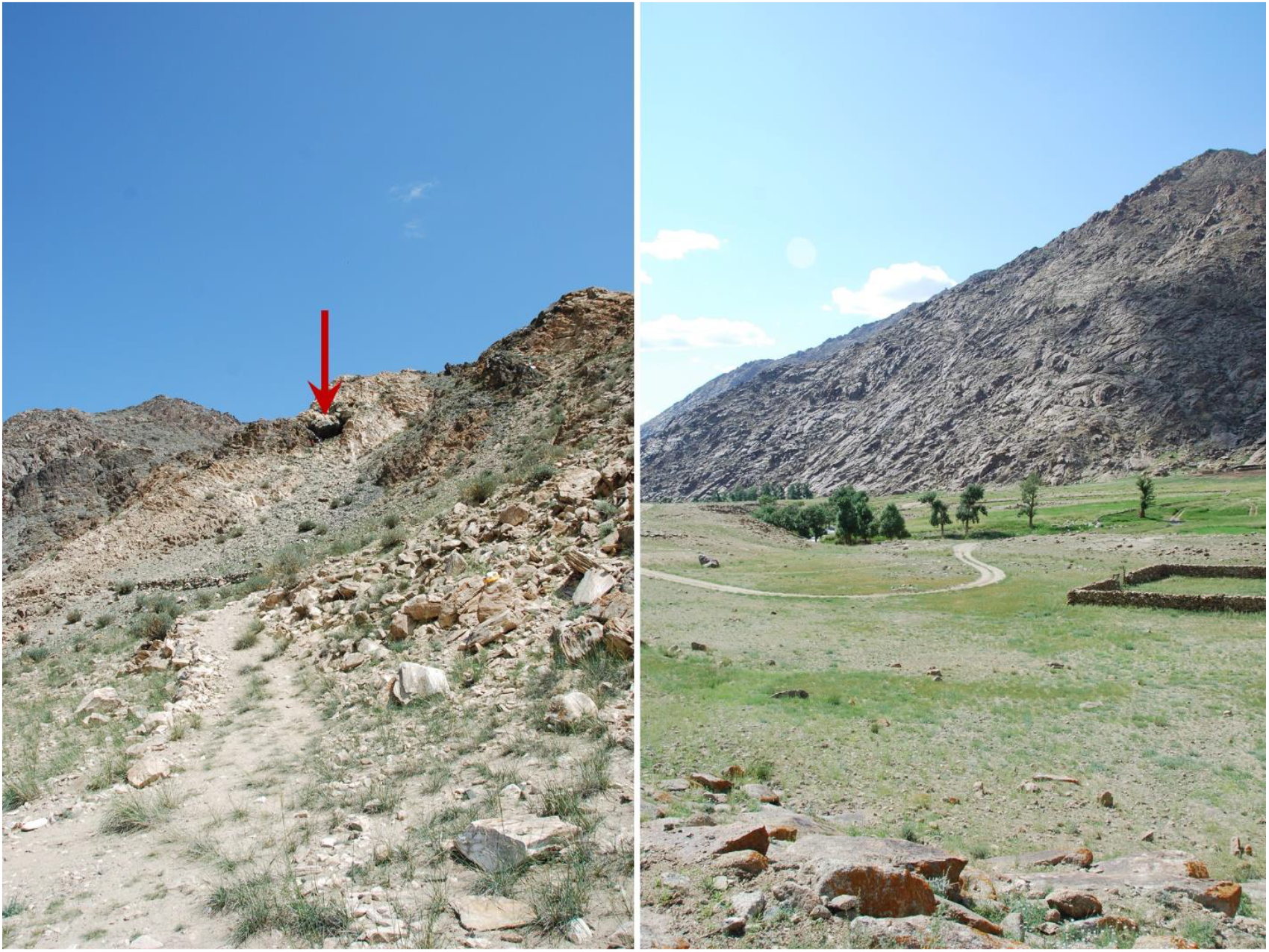
Habitat of *H. stubbei* sp. nov. from Khoya Tsenkheriyn Aguy cave (left) and Gozgor Ulaan Uul (right). At the Khoya Tsenkheriyn Aguy cave, a potential hibernation roosts (marked with an arrow), *H. stubbei* sp. nov. was captured together with *Eptesicus gobiensis, Plecotus kozlovi, Vespertilio murinus* and *Myotis davidii* on the 18/07/2011. A juvenil *H. stubbei* sp. nov. was captured at the 22/07/2011 together with *Myotis blythii* (the first record for Mongolia according to our knowledge) and *Myotis davidii* along a small stream fringed with poplar trees at the Gozgor Ulaan uul near the Bulgan gol. Photos: D. STEINHAUSER.

## Acknowledgments

We thank Bernd Gärtner, Davaa Lkhagvasuren, Jargalsaikhan Ariunbold, Idertsogt Bolorchimeg, and Klaus Thiele who contributed to the capturing of bats, collection of sample material and provided photos. We are particularly grateful to Christan Dietz for providing detailed *Hypsuogo* photos and to Hermann Ansorge for his general support. We would like to especially thank Diana Jeschke from the Natural History Museum in Görlitz for preparing some animals, as well as Irina Rusch and Katrin Krohmann from the Natural History Museum Frankfurt / Main for their support in evaluating some of the *H. savii* preparations. We acknowledge Stephan Krüger and Marie-Caroline Steinhauser for their critical reading of this manuscript.

